# Identification of secretory autophagy as a novel mechanism modulating activity-induced synaptic remodeling

**DOI:** 10.1101/2023.10.06.560931

**Authors:** Yen-Ching Chang, Yuan Gao, Joo Yeun Lee, Jennifer Langen, Karen T. Chang

## Abstract

The ability of neurons to rapidly remodel their synaptic structure and strength in response to neuronal activity is highly conserved across species and crucial for complex brain functions. However, mechanisms required to elicit and coordinate the acute, activity-dependent structural changes across synapses are not well understood. Here, using an RNAi screen in *Drosophila* against genes affecting nervous system functions in humans, we uncouple cellular processes important for synaptic plasticity from synapse development. We find mutations associated with neurodegenerative and mental health disorders are 2-times more likely to affect activity-induced synaptic remodeling than synapse development. We further demonstrate that neuronal activity stimulates autophagy activation but diminishes degradative autophagy, thereby driving the pathway towards autophagy-based secretion. Presynaptic knockdown of Snap29, Sec22, or Rab8, proteins implicated in the secretory autophagy pathway, is sufficient to abolish activity-induced synaptic remodeling. This study uncovers secretory autophagy as a novel trans-synaptic signaling mechanism modulating structural plasticity.

## INTRODUCTION

A robust functional neural circuit not only relies on proper synaptic formation and connections during development but also on the ability of the established synapses to rapidly remodel their structure and function in response to varying stimuli (*1–3*). Such activity-induced synaptic remodeling has been observed in diverse organisms, and shown to be important for behavior, cognition, and learning (*2–4*). Consequently, defective synaptic plasticity has been linked to numerous neurological disorders (*5, 6*). Studies have revealed that synaptic development and synaptic plasticity are intertwined (*7–9*), therefore making it particularly challenging to identify the molecular and cellular pathways required for activity-induced synaptic remodeling without complications from abnormal synapse development.

The *Drosophila* larval neuromuscular junction (NMJ) is as a premier model for investigating mechanisms regulating synaptic plasticity. *Drosophila* is a highly tractable genetic model organism, with approximately 75% of the known disease-causing genes conserved between fly and human (*10*). The fly NMJ is a glutamatergic synapse with stereotypical morphology, and shares similar basic molecular, physiological, and neurological properties with mammalian synapses (*11, 12*). Neuronal activity has been shown to trigger activity-induced synaptic modifications, exemplified in the fly NMJ by the formation of new synaptic boutons, enlargement of existing boutons, and increase in the abundance of postsynaptic glutamate receptors (GluR) (*13–16*). Various signaling pathways including activation of cAMP, PKA, bone morphogenic signaling, cell adhesion, and wingless pathways have been shown to regulate activity-dependent new bouton formation at the fly NMJ (*13, 14, 16–19*); however, aside from the requirement for integrin receptor activation (*15*), trans-synaptic mechanisms responsible for coordinating synaptic remodeling at established synapses (those contributing to bouton enlargement and increase in GluR abundance) remain poorly understood.

In this work, we conducted an RNAi-based genetic screen to identify the molecular pathways important for activity-induced synaptic remodeling. We made the unexpected discovery that bifurcation in the macroautophagy pathway (hereafter referred to as autophagy) differentially controls synapse development and synaptic plasticity. Although autophagy is traditionally considered a degradative pathway (*20, 21*), we show that neuronal activity both activates autophagy and suppresses degradative autophagy, favoring release through the secretory autophagy pathway. This study reveals that stimulation-induced secretory autophagy represents a novel mechanism for rapidly coordinating changes across the synapses at the fly NMJ.

## RESULTS

### An RNAi-based genetic screen to elucidate molecular mechanisms regulating activity-induced structural plasticity

To probe novel molecular pathways regulating structural plasticity, or more specifically activity-induced increase in bouton size and GluR abundance, we performed an RNAi-based genetic screen in *Drosophila*. We took advantage of the HuDis-TRiP fly RNAi collection, which contains 92% coverage of the highest-confidence fly ortholog of human disease genes (*22*). Given that altered synaptic plasticity is associated with numerous neurological disorders (*5, 6*), we focused the screen on genes affecting nervous system functions in human when mutated. We examined genes that differentially affect (1) synapse development and (2) activity-induced synaptic remodeling using RNAi-based approach, which allows the uncoupling of the two processes to identify genes most sensitive to perturbation and therefore important for each respective process. We screened through 440 *Drosophila* RNAi lines in the HuDis-TRiP collection available at the Bloomington Stock center, covering 398 unique disease-associated genes in human (Data S1). To maximize potential for identifying target genes required for activity-induced structural plasticity, RNAi knockdown was performed by crossing to a dual neuronal and muscle driver, *Elav-Gal4; 24B-Gal4*. Both basal synaptic development parameters and activity-induced changes were characterized (Fig. 1A). To assay for changes in synaptic development, we analyzed basal synaptic parameters including bouton number per NMJ (bouton number normalized to muscle surface area), size of individual type Ib boutons, and intensity of GluR in unstimulated larvae. We categorized a gene as required for normal synaptic development if any of the parameters was abnormal. To study activity-induced synaptic remodeling, we took advantage of the published protocol that a 10-min high K+ stimulation can induce rapid bouton enlargement and increase in postsynaptic GluR abundance similar to electrical stimulation (*15*). We determined the fold-change in bouton size and GluR intensity following stimulation for each RNAi line, which were normalized to the unstimulated larvae of the same genotype to account for potential differences in basal GluR levels. If the RNAi line failed to display activity-induced increases in either bouton size or GluR abundance, it was classified as a gene required for activity-induced synaptic remodeling. We classified 3 gene groups that influence synaptic morphology when knocked down: Group 1 genes have selective defects in activity-induced synaptic remodeling; Group 2 genes have defects in both synaptic development and activity-induced remodeling; Group 3 genes have defects in synaptic development but not activity-induced remodeling (Fig. 1B).

**Figure 1.**
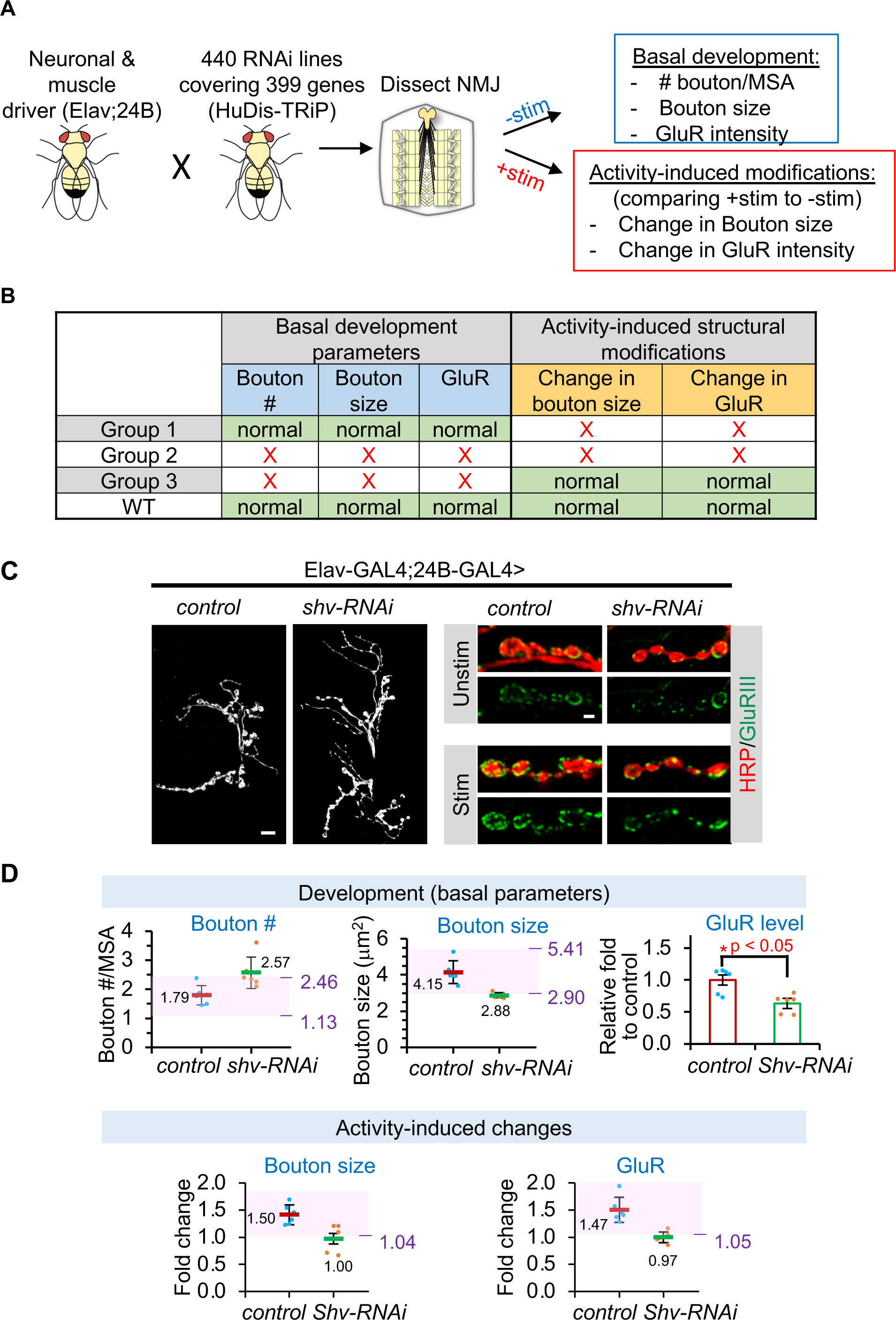
A RNAi-based genetic screen to identify genes regulating synaptic growth and activity-induced synaptic remodeling. (**A)** Summary of the RNAi-based genetic screen in *Drosophila* using the human disease TRiP (HuDis-TRiP) collection and the morphological parameters examined. **(B)** Group classification used for the genetic screen analysis. **(C)** Representative images of unstimulated (mock) and stimulated synapses at the fly neuromuscular junction (NMJ). Gray-scale images show HRP staining, which outlines the neuronal membrane, revealing a synaptic overgrowth phenotype when *shv* is knocked down using *shv-RNAi* driven by the dual neuronal and muscle *Elav-GAL4;24B-GAL4* driver. Scale bar = 10 μm. Right images show synaptic boutons stained with HRP (red) and GluRIII antibody (green). Scale bar = 2 μm. **(D)** Graphs showing that the cutoff values used in the genetic screen can reliably detect altered synaptic development and activity-induced synaptic remodeling in *shv-RNAi*. Shaded areas highlight cutoff values derived from 6 driver control NMJs. Bouton number is normalized to muscle surface area (MSA), and fold change is calculated by comparing stimulated to mock unstimulated NMJs of the same genotype. Values are mean ± S.E.M.

Due to the large number of samples and images generated from the screen, we performed an expedited analysis to assess both basal developmental parameters and activity-induced changes (see Methods). We confirmed that this expedited analysis indeed detected activity-induced structural modifications in the driver control at a level comparable to those reported previously (Fig. 1C, D), and that the established cutoff parameters successfully detected altered synapse development and activity-induced synaptic modifications in RNAi against shriveled (*shv*), a gene previously shown to regulate both processes (*15*).

The result of the genetic screen is shown in Fig. 2A. All morphological parameters examined showed normal distribution (fig. S1). There were 71 genes that did not result in any synaptic phenotype and 24 genes that caused lethality when knocked down (Data S2); these were not analyzed further. The majority of the genes showed defects in both synapse development and activity-induced synaptic remodeling (141 genes, Group 2; Data S2). We also uncovered 94 genes that displayed selective defects in activity-induced synaptic remodeling when knocked down (Group 1; Data S2) and 68 genes that disrupted synapse development but not activity-induced synaptic remodeling (Group 3; Data S2). There were some redundant RNAi lines targeting the same human disease gene in the genetic screen. While most overlaps resulted in the same group categorization, a few did not, likely due to the different extent of gene knockdown. These genes were therefore classified by its more severe phenotype, i.e. a Group 1 and wildtype result for redundant RNAi lines against the same gene would be classified as Group 1, whereas a Group 1 and Group 2 result for redundant RNAi lines would be classified as Group 2. Interestingly, we never observed disparate classification for a gene in Groups 1 and 3, which are genes specifically affecting acute synaptic remodeling and synapse development, respectively. Together, these results reveal that while synapse development and activity-induced synaptic remodeling could involve overlapping proteins and pathways, it is still possible to uncouple them, as this RNAi knockdown approach identifies key genes most sensitive to alternations.

**Figure 2.**
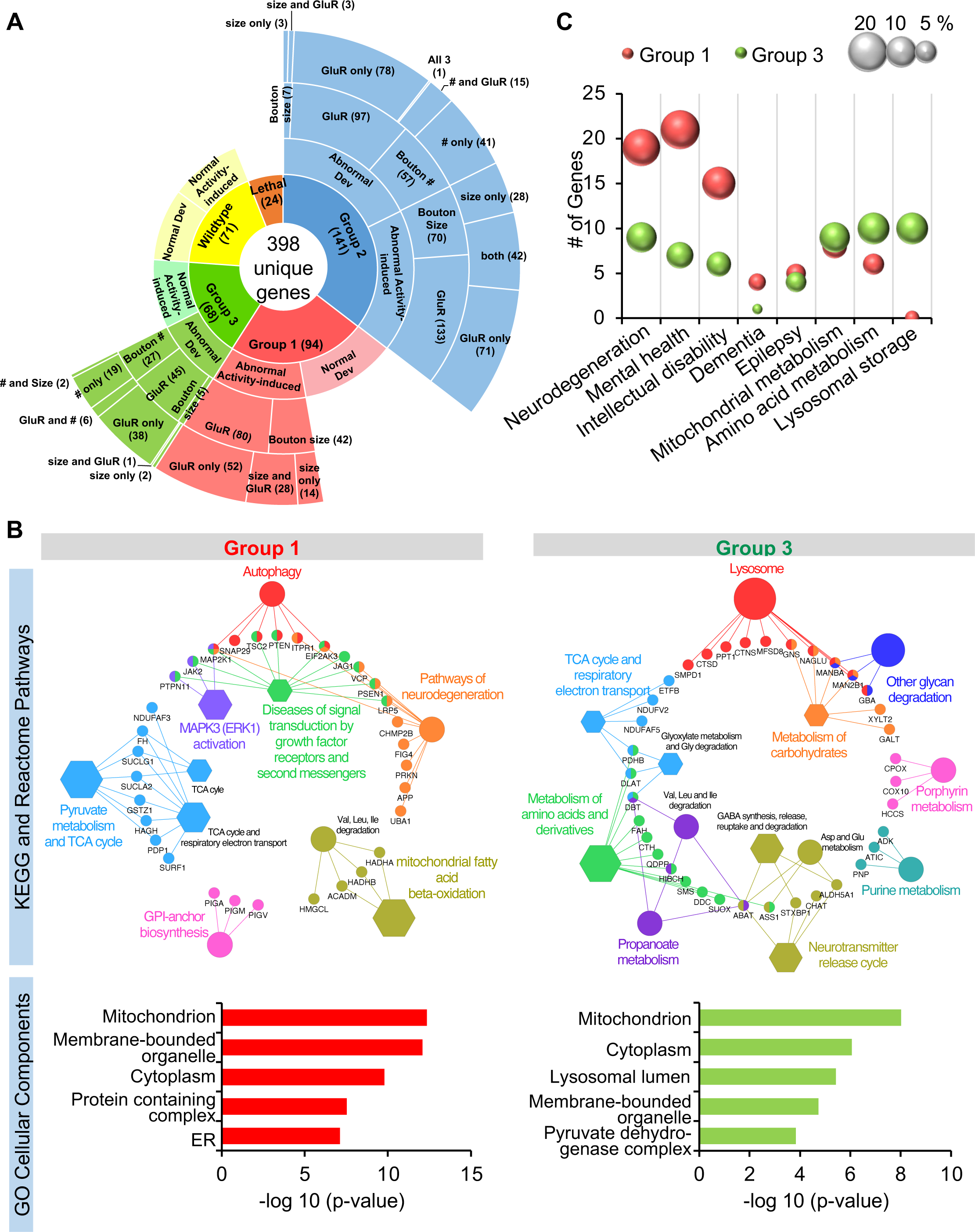
Bioinformatics analysis of genes selectively perturbing activity-induced synaptic remodeling or synaptic development in the genetic screen. **(A)** Sunburst graph summarizing the results of the genetic screen for 440 RNAi lines against 398 unique human disease orthologs. **(B)** KEGG and Reactome pathway analyses visualized using ClueGo in Cytoscape. Circles represent KEGG pathway and hexagons represent Reactome pathway terms, and larger size represent higher significance. Only term description with p-value < 0.05 are shown. Individual nodes (small circles) list the individual genes in the functional pathways. Human gene orthologs were used in bioinformatics analysis. Lower graph shows the GO cellular compartment analysis for Group 1 and 3 genes. **c**, Graphs show the number of genes identified in the respective disease categories. Size of the sphere indicates percentage of genes in each group belonging to the disease category. Analysis was performed using the DISEASE Database in the String plugin in Cytoscape.

### Mechanisms regulating synapse development and activity-induced synaptic remodeling

To obtain a broad understanding of the molecular pathways regulating synapse development, we examined all genes that showed altered basal synaptic development parameters (Groups 2 and 3 genes). KEGG and Reactome functional enrichment analyses highlight that metabolism pathway, along with lipid, amino acid, and energy metabolism modulate synaptic bouton number (fig. S2a). An enrichment in genes affecting glycosylation and autophagy was also identified. We also examined genes that altered bouton size in unstimulated NMJs when knocked down. Due to the small number of genes uncovered from the screen (11 genes), no functional enrichment was found using KEGG and Reactome analyses, but manual curation identified the presence of 3 mitochondrial proteins and 1 peroxisomal proteins, implying that lipid and energy metabolism could be important for the regulation of bouton size. Lastly, we examined RNAi lines with altered basal GluR levels. We found metabolism, lysosomal function, breakdown of glycosaminoglycan, and glycolysis pathways are important to maintain basal GluR levels (fig. S2B), highlighting that proteostasis and energy metabolism are particularly important.

Next, we aimed to identify key signaling pathways necessary for the rapid, activity-dependent synaptic changes by comparing genes selectively disrupting activity-induced synaptic remodeling (Group 1; Data S2) versus those that affect development (Group 3; Data S2). These two disparate groups allow us to independently examine key proteins affecting synapse development or synaptic plasticity when perturbed. Interestingly, KEGG and Reactome functional pathway analyses revealed that Group 1 showed an enrichment in genes associated with neurodegeneration, as well as autophagy and protein processing in the ER, consistent with an enrichment of ER proteins determined using the GO Cellular Component enrichment analysis (Fig. 2B). Genes selectively disrupting synapse development (Group 3) are highly enriched in lysosomal pathway and amino acid metabolism. GO Cellular Component enrichment analysis also showed a comparable increase in lysosome distribution. We also used the DISEASES database to investigate the relationship between group classification and human diseases. We found that neurological disorders, mental health disorders, intellectual disability, and dementia showed a 2-fold enrichment in genes affecting activity-induced synaptic remodeling than synapse development (Fig. 2C and fig. S3). Epilepsy and metabolic disorders showed approximately equal distribution, whereas amino acid metabolism disorder and lysosomal storage disease displayed significant enrichment in genes affecting synapse development. These findings reaffirm that structural plasticity is important for cognition and memory.

**Figure 3.**
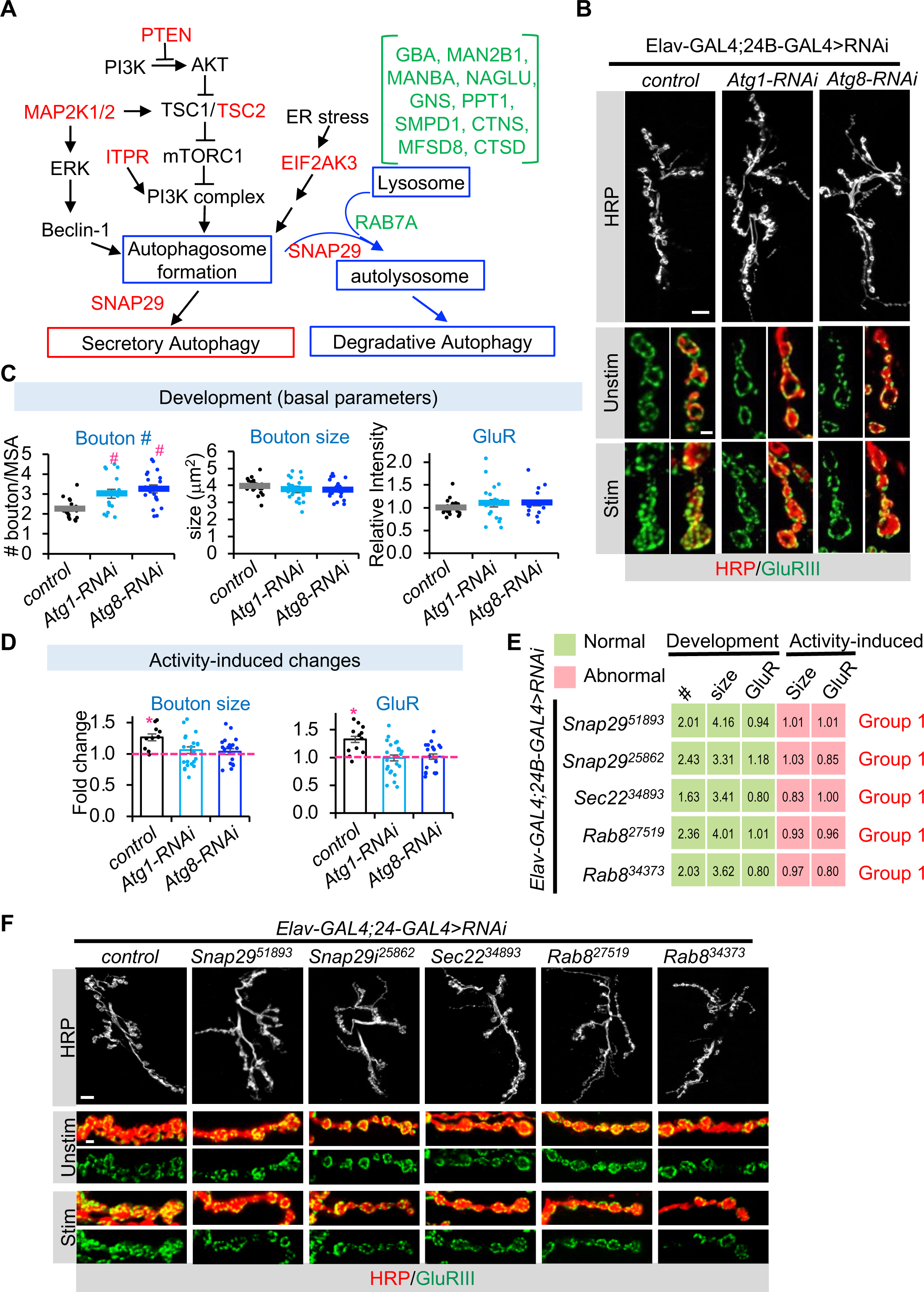
Bifurcation in autophagy pathway differentially regulates activity-induced synaptic remodeling and synaptic development. **(A)** Autophagy pathway with red text highlighting Group 1 genes and green text highlighting Group 3 genes identified in the genetic screen. Blue outline highlights conventional degradative autophagy pathway. **(B)** Perturbing autophagy core molecular components caused synaptic overgrowth and defective synaptic plasticity. Scale bar = 10 μm for gray-scale images, and 2 μm for colored images. Quantification of (**C**) basal synaptic development parameters and **(D)** activity-induced synaptic remodeling for the indicated fly lines. Bouton number is normalized to muscle surface area (MSA), and fold change is calculated by comparing stimulated to mock unstimulated NMJs of the same genotype. All values are mean ± s.e.m. One-way Anova followed by Tukey’s multiple comparison test was used to compare between control and unstimulated samples across genotypes. Student’s t-test was used to compare between unstimulated and stimulated NMJs of the same genotype. # p ≤ 0.05 when compared to unstimulated control. *p ≤ 0.05 when comparing stimulated to unstimulated NMJs. **(E)** Knockdown of proteins involved in secretory autophagy release pathway in both muscles and neurons selectively blocked activity-induced synaptic remodeling. Same cutoff values from the genetic screen were used for group classification. **(F)** Gray-scale images of unstimulated NMJs labeled with HRP (top row). Scale bar = 10 μm. Colored images show unstimulated and stimulated synaptic terminals stained with HRP (red) and GluRIII (green). Scale bar = 2 μm.

### Role of autophagy in activity-induced synaptic remodeling

The finding that autophagy regulates activity-induced synaptic remodeling while lysosomal functions are important for synapse development seemed paradoxical, since canonical autophagy pathway leads to degradation by lysosomes (*21*). We therefore performed a detailed pathway analyses for Groups 1 and 3 genes identified in the KEGG pathway. We found that genes selectively affecting activity-induced synaptic remodeling included those that modulated autophagy activation rather than required for the degradative step of autophagy (Fig. 3A; red text). It also contained Snap29, a protein involved in multiple membrane fusion steps including autophagosomes-lysosome fusion and autophagosome fusion with the plasma membrane for release of autophagosomes during secretory autophagy (*23–25*) (Fig. 3A). Conversely, RNAi lines that selectively altered synapse development identified in the screen (Group 3) included 10 different lysosomal degradative enzymes or lysosomal membrane proteins (green text ; Fig. 3A), as well as RAB7, a protein important for autophagosome-lysosome fusion and therefore degradative autophagy (*26, 27*). These findings imply that activity-induced synaptic remodeling does not require degradative autophagy but is most sensitive to disruptions in autophagy-dependent events prior to lysosomal degradation.

To further understand the involvement of autophagy in synaptic plasticity, we first tested the effects of perturbing proteins in the core autophagy machinery, or proteins directly required for autophagosome formation. RNAi against autophagy-related proteins atg1 or atg8, which should disrupt both autophagosome formation and subsequent degradative autophagy (*28–30*), altered synaptic growth and impaired activity-induced remodeling (Fig. 3B-D). These data confirm that autophagy in general is important for both processes and are consistent with our genetic screen results. They also support that idea that bifurcation in the conventional degradative autophagy pathway differentially regulate activity-induced synaptic remodeling and development.

Aside from degradation by lysosomes through the conventional autophagy pathway, autophagosome can also fuse directly with the plasma membrane to release its contents extracellularly through secretory autophagy pathway (*31*) (Fig. 3A). While molecular mechanisms regulating this unconventional secretory autophagy pathway is poorly understood, Snap29, Sec22b, and Rab8a have all been implicated in this pathway by driving autophagosome fusion with the cell membrane for secretion (*25, 32–34*). We have already identified Snap29 as a protein selectively required for activity-induced synaptic remodeling (Group 1 gene) and have now confirmed the result using a second RNAi for Snap29 (*Snap-RNAi^25862^*) (Fig. 3E, F). We also examined the effects of knocking down Sec22 and Rab8, which are fly orthologs of human Sec22b and Rab8a, respectively. Sec22b is a SNARE protein that collaborates with Snap 29 to regulate secretory autophagy (*25, 33*), and both Sec22b and Rab8a can facilitate the release of interleukin through secretory autophagy pathway in mammalian cells (*25, 32*). We found expression of Sec22-RNAi or Rab8-RNAi lines using the dual neuron and muscle driver disrupted activity-induced synaptic remodeling but not synaptic development (Fig. 3E, F). These results are consistent with the involvement of unconventional secretory autophagy pathway in regulating acute, activity-induced synaptic changes.

### Presynaptic knockdown of secretory autophagy molecular machinery is sufficient to block activity-induced synaptic bouton enlargement and increase in GluR abundance

We speculated that secretory autophagy activation during neuronal activity serves as a paracrine signaling mechanism that allows communication and coordination of synaptic changes across the synapse. To delineate the spatial requirement for secretory autophagy in activity-induced synaptic remodeling, we independently knocked-down Snap29, Rab8, and Sec22 in neurons using Elav-GAL4 or muscles using 24B-GAL4 driver. We found that presynaptic knockdown of secretory autophagy molecular components is sufficient to block activity-induced synaptic remodeling (Fig. 4), while knockdown in muscles resulted in a profound, partial change in synaptic phenotypes (fig. S4). Specifically, *Rab8-RNAi* expression in muscles could still undergo activity-induced bouton size enlargement, whereas Sec22-RNAi in muscles reduced GluR intensity during development. Sec22 has multiple functional roles including regulation of protein trafficking (*35*), suggesting that Sec22 may regulate the trafficking of GluR in muscles. Together, these data imply that although both muscles and neurons can utilize autophagy-based secretory pathway to non-cell autonomously regulate synaptic bouton size and GluR abundance, communication between the pre- and postsynapse may influence the overall phenotype, and that proteins regulating secretory autophagy may have different dominant functional roles in distinct cell types. Nonetheless, our results indicate that neuronal secretory autophagy pathway is sufficient to promote synaptic bouton size enlargement and coordinate changes in postsynaptic GluR levels in response to stimulation.

**Figure 4.**
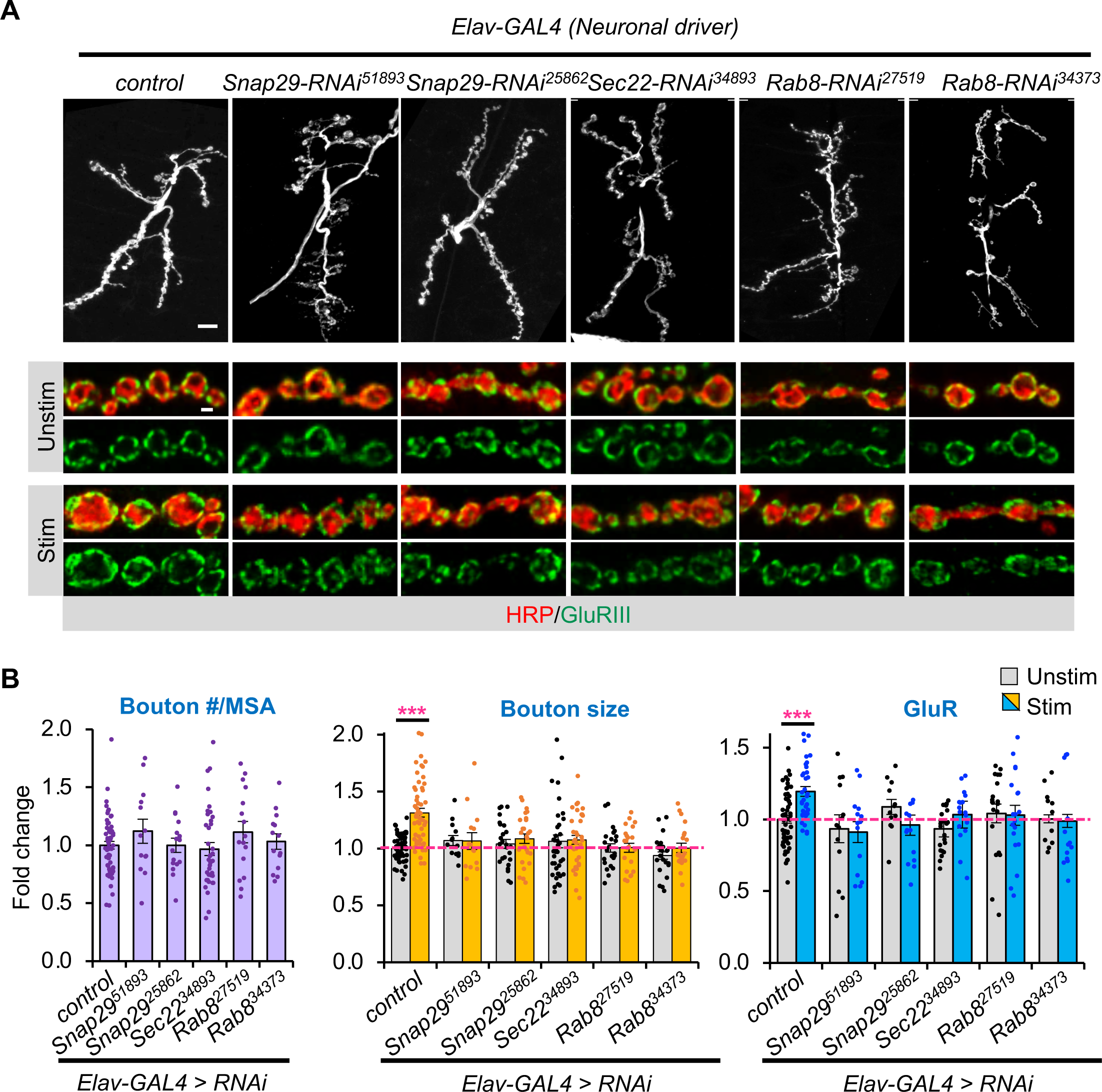
Neuronal knockdown of secretory autophagy molecular components. **(A)** Gray-scale images of unstimulated NMJs labeled with HRP (top row). Scale bar = 10 μm. Colored images show unstimulated and stimulated synaptic terminals stained with HRP (red) and GluRIII (green). Scale bar = 2 μm. **(B)** Quantification of fold change in bouton number (normalized to muscle surface area), bouton size, and GluR intensity as indicated. All values are mean ± s.e.m. Statistics: One-way Anova followed by Tukey’s multiple comparison test was used to compare between control and unstimulated samples across genotypes. Student’s t-test was used to compare between unstimulated and stimulated NMJs of the same genotype. *** p ≤ 0.001 when comparing stimulated to unstimulated NMJs.

### Stimulation blocks autophagosome-lysosome fusion but enhances secretory autophagy

Given that disrupting autophagy-based secretory pathway in neurons is sufficient to block synaptic remodeling, we next set out to monitor autophagy during plasticity-inducing stimulation in neurons. Autophagy is a highly dynamic process that involves autophagosome formation, fusion of autophagosomes with lysosomes to form autolysosomes, and the turnover and removal of autolysosomes (*20, 21*) (Fig. 5A). We monitored total autophagic vesicles using mCherry-atg8a, as the pH independent nature of mCherry allows labeling of both autophagosomes and autolysosomes (*36*). Lysosomes were monitored using the marker Lamp1-GFP, and overlap between mCherry-Atg8a and Lamp1-GFP represent the autolysosomes (*36*) (Fig. 5A). We found that stimulation did not significantly alter the intensity nor the number of mCherry-Atg8a and Lamp1-GFP spots at the NMJ, but it significantly reduced the number of autolysosomes as determined by mCherry-Atg8a and Lamp1-GFP signal colocalization (Fig. 5B-E; fig. S5A,B). This stimulation-dependent decrease in autolysosomes could either be due to increased autophagy flux or a block in autophagosome-lysosome fusion. To discern these two cases, we monitored autolysosome number by further depleting autophagosome-lysosome fusion using Rab7-RNAi. We expected that blocking autolysosome formation will exhaust the existing autolysosomes in an activity-dependent manner if enhanced autophagic degradation is the underlying cause. However, Rab7-RNAi showed a similar extent of decrease in the number of autolysosomes post-stimulation compared to the control (Fig. 5C), revealing that neuronal activity does not enhance autophagy flux but rather inhibits autophagosome-lysosome fusion. Note that Rab7-RNAi also reduced the basal autolysosome number, consistent with its role in mediating autophagosome-lysosome fusion (*27*). To further validate this result, we visualized the early part of the autophagy pathway using GFP-Atg8a, as the pH sensitive nature of GFP permits only the labeling of autophagosomes prior to fusion with lysosomes (*36*). We observed a striking increase in GFP-atg8a intensity and the number of punctate spots following stimulation in control and Rab7-RNAi NMJs (Fig. 5F, G; fig. S5C), implying enhanced autophagy activation and consistent with stimulation-induced block in autophagosome-lysosome fusion. Note that we were not able to use the tandemly labeled mCherry-GFP-Atg8a for the above experiments, since this construct did not show sufficient signal to allow detection at the fly NMJ.

**Figure 5.**
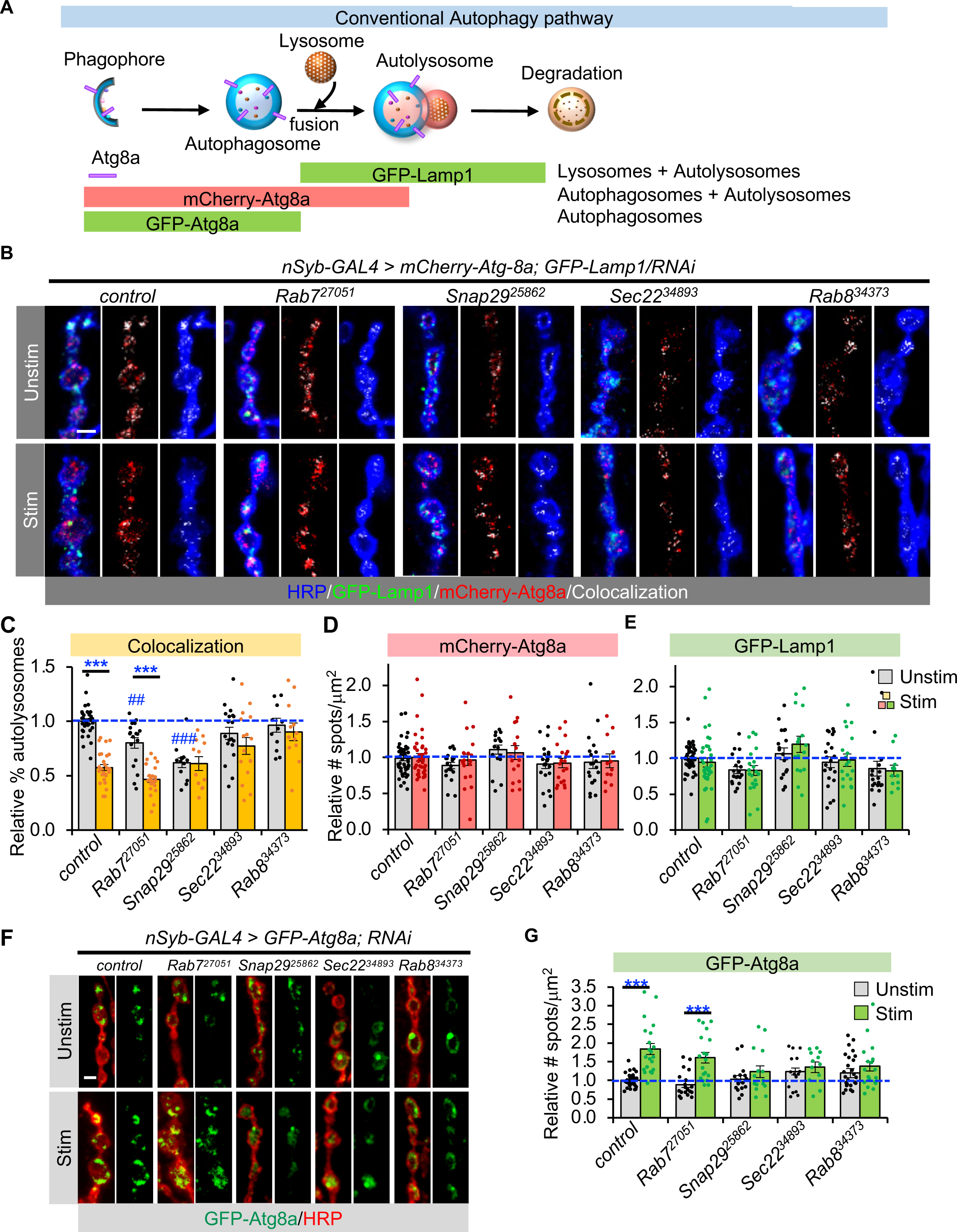
Stimulation enhances autophagosome formation while inhibiting autophagosome-lysosome fusion. **(A)** Schematic outline of conventional degradative autophagy pathway and fluorescent markers used to detect the different organelles. **(B)** Representative images showing the number of autolysosomes decreased following stimulation. Colocalization between mCherry-Atg8a (red) and GFP-Lamp1 (red) represent the autolysosomes, which are shown in white. HRP (blue) outlines the synaptic bouton membrane. **(C)** Quantification of the fold change in autolysosomes in unstimulated and stimulated NMJs. To account for differences due to the number of autophagic vesicles, the number of autolysosomes was normalized to the number of autophagosomes (mCherry-Atg8a spots) to obtain the percent of autolysosomes in each condition. Fold change is determined by comparing to the mock unstimulated control. **(D)** Fold change in the number of punctate mCherry-Atg8a signal and **(E)** GFP-Lamp1 signal in unstimulated and stimulated NMJs normalized to bouton area. **(F)** Representative images of GFP-Atg8a. Stimulation increased GFP-Atg8a signal, indicating an increase in autophagy initiation. **(G)** Fold change in GFP-Atg8a punctate spots in unstimulated and stimulated NMJs for the indicated genotypes. Scale bar = 2 μm in (**B**) and (**F**). All panels show mean ± s.e.m. Statistics: (**C**) and (**E**) One-way Anova followed by Tukey’s multiple comparison test was used to compare between control and unstimulated samples across genotypes. Student’s t-test was used to compare between unstimulated and stimulated NMJs of the same genotype. ## p ≤ 0.01; ### p ≤ 0.001 when comparing unstimulated samples of different genotypes to control. *** p ≤ 0.001 when comparing stimulated to unstimulated NMJs.

Next, we monitored autophagy in Snap29, Sec22, and Rab8-RNAi lines. Knockdown of proteins important for secretory autophagy shunted the pathway towards degradation, since the number of autolysosomes (mCherry-Atg8a and Lamp1-GFP colocalization) no longer declined and autophagosomes (GFP-Atg8a) no longer increased post-stimulation (Fig. 5B, C, F, G). Interestingly, Snap29 also displayed reduced autolysosome signal even without stimulation, consistent with reports that Snap29 also mediates autophagosome-lysosome fusion (*23, 24*). Together, our results imply that neuronal activity promotes autophagy activation but inhibits autophagosome-lysosome fusion, whereas disrupting autophagy-based secretory pathway drives the pathway towards degradative autophagy pathway.

What is the physiological significance of this stimulation-induced pause in autophagosome-lysosome fusion? We propose that a block in autolysosome formation directs the pathway towards secretory autophagy. Currently there are no known markers for unconventional autophagy-based secretory pathways in neurons. Based on our genetic screen results that knockdown of one of the lysozyme genes, lysozyme P (LysP), resulted in a selective defect in activity-induced synaptic remodeling (Data 2) and a report that lysozyme could be released through secretory autophagy pathways by Paneth cells upon bacterial invasion (*37*), we explored the possibility that lysozyme is released via this route at the fly NMJ upon stimulation. First, we measured the levels of extracellular lysozyme using a non-permeabilizing staining condition. Stimulation substantially elevated the presence of extracellular lysozyme at the synapse (Fig. 6A). To differentiate the source of extracellular lysozyme, we independently knocked down LysP in neurons or in muscles. Neuronal expression of LysP-RNAi substantially lowered extracellular lysozyme level in unstimulated NMJs while muscle knockdown only mildly suppressed lysozyme level (Fig. 6B). Furthermore, neuronal knockdown of LysP abolished the activity-induced increase in extracellular lysozyme signal at the synapse while muscle knockdown did not (Fig. 6A, B). Together, these data indicate that neurons can release lysozymes to the extracellular milieu upon plasticity-inducing stimulation.

**Figure 6.**
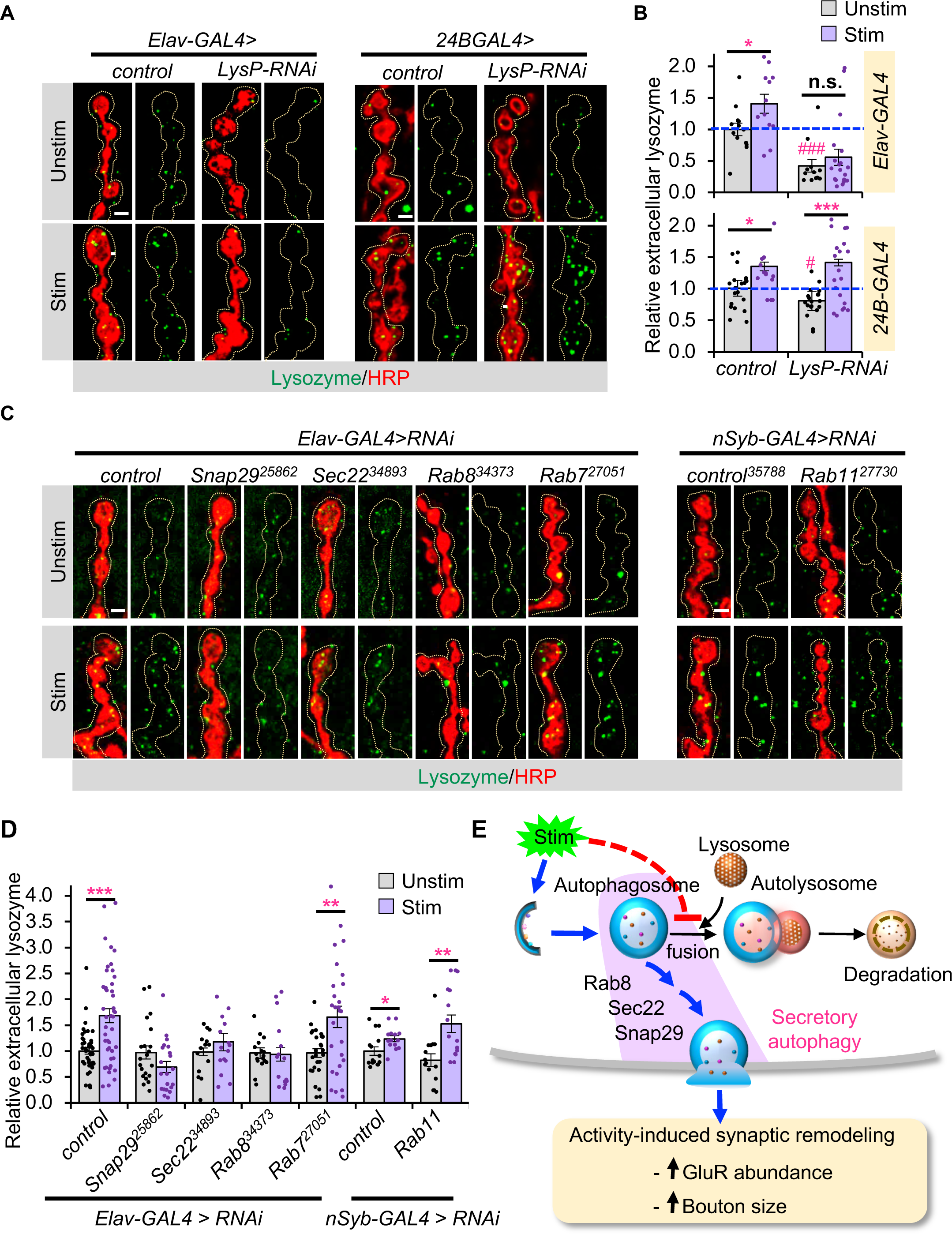
Activation of secretory autophagy by neuronal activity and model for autophagy-based regulation of synaptic plasticity. **(A)** Extracellular lysozyme staining performed using detergent-free condition. **(B)** Neuronal knockdown of LysP-RNAi diminished activity-induced extracellular lysozyme staining at the synapse while muscle knockdown did not. Scale bar = 2 μm. **(C)** Representative images of extracellular lysozyme staining in the RNAi lines as indicated. For **(A)** and **(C)**, dashed outline shows the area used to determine extracellular lysozyme level, which is 0.5 μm beyond the neuronal membrane. Scale bar = 2 μm. **(D)** Quantification of extracellular synaptic lysozyme staining. Blocking secretory autophagy abolished activity-induced increase in extracellular lysozyme staining. All values are mean ± s.e.m. Statistics: (**B**) and (**D**) One-way Anova followed by Tukey’s multiple comparison test was used to compare between control and unstimulated samples across genotypes. Student’s t-test was used to compare between unstimulated and stimulated NMJs of the same genotype. # p ≤ 0.05; ### p ≤ 0.001 when comparing unstimulated samples of different genotypes to control. * p ≤ 0.05; ** p ≤ 0.01; *** p ≤ 0.001 when comparing stimulated to unstimulated NMJs of the same genotype

To determine whether synaptic lysozyme is released via autophagy-based release pathway, we examined lysozyme release while disrupting proteins involved in secretory autophagy. Neuronal knockdown of Snap29, Rab8, and Sec22 each blocked the activity-induced increase in extracellular lysozyme (Fig. 6C,D), suggesting that neuronal activity activates secretory autophagy. We also tested the involvement of the exosome release pathway in lysozyme secretion. To this end, we knocked down the Rab11 GTPase in neurons, which has been shown to modulate endosome recycling and exosome release at the fly NMJ (*38*). Expression of *Rab11-RNAi* using the Elav-Gal4 driver caused lethality, we thus used another pan-neuronal driver, *nSyb-GAL4*, which resulted in viable progeny perhaps due to the lower expression level. Neuronal knockdown of Rab11 did not affect stimulation-induced increase in extracellular lysozyme (Fig. 6C, D). To ensure that Rab11-RNAi efficiently altered exosome release, we monitored the levels of extracellular neuroglian (Nrg), a protein known to be release in extracellular vesicles through the exosomal pathway (*38*). Indeed, *Rab11-RNAi* expression in neurons reduced extracellular Nrg levels that were not further elevated by neuronal activity (fig. S6A, B). Conversely, Snap29, Rab8, or Sec22 knockdown displayed normal activity-dependent increase in Nrg, confirming the release of lysozyme and Nrg by independent pathways. We also found that Rab11-RNAi affected both synapse development and activity-induced synaptic remodeling (fig. S6C, D). Interestingly, *Rab7-RNAi*, a protein mediating both autophagosome-lysosome fusion and endolysosomal fusion (*39*), increased the basal levels of extracellular Nrg (fig. S6A, B), further implying that Nrg may be under homeostatic regulation by lysosomes. Taken together, these data reveal (1) lysozyme can be used to monitor secretory autophagy pathway and is distinct from exosomal release pathway, (2) exosome release pathway can also modulate synapse development and activity-induced synaptic remodeling, and (3) neuronal activity activates secretory autophagy.

## Discussion

Using an RNAi screen against fly orthologs of human disease genes, we discovered that autophagy-based secretory pathway is a novel mechanism regulating activity-induced synaptic remodeling at the fly NMJ. While autophagy has previously been linked to synaptic plasticity, it is thought to act through the degradative autophagy pathway to maintain protein homeostasis (*40*). Our data supports a new model in which neuronal activity promotes autophagy activation but dampens autophagosome-lysosome fusion, thereby driving the pathway towards secretory autophagy release. This results in an activity-dependent trans-synaptic signal that enables rapid communication and coordination of synaptic changes across the synapse (Fig. 6E).

The RNAi genetic screen presented in this paper focused on genes associated with neurological disorders in human. Albeit this is a biased screen, this approach not only allows us to compile information on synaptic changes associated with various diseases of the nervous system, but also to quickly survey candidate genes to identify genes most sensitive to changes in levels and therefore most important for synapse development and synaptic plasticity. Consistent with the idea that synaptic plasticity is important for cognition and memory, we found that intellectual disability, neurodegenerative and mental health disorders are more likely to associate with defects in activity-induced synaptic remodeling than synapse development. We acknowledge that while this RNAi-based approach does not conclusively eliminate the involvement of a gene when the result is negative, it is advantageous as it bypasses problems commonly associated with knockout of crucial genes, such as lethality.

Autophagy is classically known as a degradative process in which autophagosomes fuse with lysosomes to breakdown its contents to maintain proteostasis (*20, 21*). Results showing that knockdown of 10 different lysosomal proteins did not affect activity-induced synaptic remodeling but altered synapse development strongly support that acute, activity-dependent synaptic changes does not rely on degradative autophagy. Aside from the conventional degradative pathway, autophagosomes have recently been shown to be released in response to infection or stress through a processed called secretory autophagy (*31*). Our data strongly support that neurons utilize secretory autophagy as a mechanism to communicate and coordinate activity-induced synaptic remodeling. First, consistent with reports that neuronal activity induces autophagy activation (*41–43*), we observed elevated GFP-Atg8a signal post-stimulation (Fig. 5F, G). Second, knockdown of proteins implicated in secretory autophagy pathway in neurons is sufficient to block activity-induced bouton enlargement and postsynaptic GluR abundance (Fig. 4). Although Snap29, Rab8, or Sec22 each play multiple roles in regulating vesicle trafficking and membrane fusion, they overlap in their ability to facilitate secretory autophagy (*23–25, 33–35*). Third, we identified that lysozyme is released through secretory autophagy pathway, and present evidence that neuronal activity enhances lysozyme release (Fig. 6). Lysozyme is typically known as an antimicrobial substance (*44*), but it also has other less appreciated functions including regulation of bone development and activation of toll-like receptors to mediate neuropathic pain (*45, 46*). Interestingly, toll-like receptor signaling can modulate multiple processes including neurogenesis and synaptic plasticity (*47, 48*). Our genetic screen also revealed that knockdown of LysP abolished activity-induced synaptic remodeling. It will be interesting to delineate mechanisms by which lysozyme signaling regulate synaptic plasticity in the future. Note that while we observed an increase in GFP-Atg8a signal consistent with increased autophagy activation during neuronal activity, we did not detect an increase in the number of mCherry-Atg8a spots (which also labels autolysosomes in addition to autophagosomes) as reported previously for prolonged neuronal stimulation (*41, 43*). This difference is likely due to different stimulation conditions, as a 30-minute stimulation was required to elevate mCherry-Atg8a signal (*41*). This suggests that prolonged stimulation may ultimately activate a different autophagy program to maintain synaptic homeostasis. Future studies examining autophagy regulation during varying neuronal activity will shed light on how neurons utilize different autophagic pathways to optimize function.

We would like to note that while this study focused on secretory autophagy as a mechanism regulating activity-induced synaptic remodeling, we do not rule out the possibility that crosstalk between endosomal and autophagy release pathways also contribute. In fact, we found that knockdown of Rab27A, a protein mediating release of extracellular vesicles downstream of amphisomes (formed by endosome-autophagosome fusion) (*49, 50*), in our genetic screen as a protein affecting activity-induced remodeling. Rab8a has also been shown to play dual roles in regulating secretory autophagy and extracellular vesicle release downstream of endosome-autophagosome pathway (*50*). Thus, future experiments examining crosstalk between endosome, autophagosome, and lysosome pathways will provide insights into how neurons integrate various cellular pathways to coordinate activity-dependent synaptic changes. Identification of cargos released through autophagy-based secretion will also facilitate our understandings of the physiological significance of this novel signaling pathway and its role in neurodegenerative disorders.

### Materials and Methods Drosophila culture and stocks

Flies were grown on standard cornmeal, yeast, sugar and agar medium at 25 °C under a 12 hours dark/light cycle. The TRiP RNAi lines used in the genetic screen were obtained from the Bloomington Drosophila Stock Center (BDSC) and listed in Supplmenentary Data S1. RNAi stocks from the Hu-Dis TRiP collection were included in the screen if “nervous system disease” was listed in the disease description. The following stocks were also obtained from the BDSC (stock number in parenthesis): UAS-Atg8a-RNAi (#58309), UAS-Atg1-RNAi (#44034), UAS-Snap29-RNAi (#25862), UAS-Rab8a-RNAi (#34373, #27519), UAS-Sec22-RNAi (#34893), UAS-Rab11-RNAi (#27730), UAS-mCherry-Atg8a (#37750), UAS-GFP-Atg8a (#52005), UAS-GFP-Lamp1;nSyb-GAL4 (#42714), *Elav-Gal4* (#458), and 24B-GAL4 (#1767), *nSyb-GAL4* (gift from Dr. Simpson), *UAS-LUC-Valium10* (TRiP RNAi control vector; #35788). All standard balancer stocks were obtained from Bloomington Stock Center.

### Dissection and stimulation

Third-instar larval body wall were dissected in normal HL-3 solution without Ca^2+^ (NaCl 110mM, KCl 5mM, MgCl_2_ 10mM, sucrose 30mM, HEPES 5mM, EGTA 1mM, trehalose 5mM, NaO_3_ 10mM, pH 7.2), leaving the brain and peripheral nerves intact. High K^+^ stimulation was performed by incubating the dissected NMJ preps with 90mM KCl in HL-3 solution with Ca^2+^ (NaCl 25mM, KCl 90mM, MgCl_2_ 10mM, CaCl_2_ 1.5mM, sucrose 30mM, HEPES 5mM, trehalose 5mM, NaHCO_3_ 10mM, pH 7.2) for 10 min, and then placed in resting solution for 2 minutes (normal HL-3 solution without EGTA). Mock, unstimulated samples were included in each experiment, except instead of high KCl solution, they were left in the resting buffer for 10 minutes and processed the same way. Samples were then fixed with either bouin’s fixatives for 3 mins (if performing GluR staining) or with 4% paraformaldehyde solution for 25 minutes at room temperature.

### Immunocytochemistry

Fixed samples were washed with 0.1% triton X-100 in PBS (PBST) or with PBS for detergent-free condition. Samples were blocked with 5% normal goat serum in PBST or PBS as indicated. Antibodies were diluted in blocking solution and used as following: rabbit anti-GluRIII, 1:1000 (generated in this study); mouse anti-lysozyme, 1:200 (Biorad SB1), mouse anti-neuroglian, 1:100 (DSHB BP104), Alexa-647 and Cy3-conjugated anti-HRP, 1:100 (Jackson ImmunoResearch). Secondary antibodies used were Alexa-488 or 405 conjugated, 1:250 (Invitrogen). The rabbit anti-GluRIII antibody was generated by Pocono Rabbit Farms (Canadensis, PA), using the 22 C-terminal residues of GluRIII (-QGSGSSSGSNNAGRGEKEARV) as the immunogen. The serum was affinity purified and used at 1:1000 dilution.

### Imaging

Images of synaptic boutons from muscle 6/7, A2 or A3 were acquired using either a Zeiss LSM800 or Olympus FV3000 confocal microscope. To expedite image capture for the genetic screen, 3-z stacks at 2 μm interval were used to capture the entire depth of the synaptic boutons at 40x (Zeiss) or 4-z stacks at 1.5 μm interval were captured using Olympus at 60x. A single plane 10x image was also taken for each animal in order to determine the muscle surface area. To establish changes in basal GluR level, unstimulated, mock control larvae were always dissected and imaged in parallel with test genotypes for each experiment. Stimulated and unstimulated animals per genotype were also performed and imaged in parallel using the same conditions in order to measure fold-change in bouton size and GluR intensity. A minimum of 5 NMJs from a minimum of 2 larvae were imaged per condition and per genotype for the genetic screen. For all other experiments, images were acquired using an Olympus FV3000 confocal microscope at 60x with 1.6 zoom.

### Image analysis

For the genetic screen, an expedited analysis using 15 type Ib boutons belonging to a minimum of 2 branches of axons at the M6/7 were manually circled in Image J to determine the average bouton size and average GluR intensity. Control NMJ (w1118 crossed to driver control) was used to establish the cutoff parameters. For morphological parameters that could be quantified by absolute values such as bouton number and size, and the fold-change in bouton size and GluR intensity following stimulation (normalized to unstimulated NMJ of the same genotype), a value that is 2 standard deviations (95% confidence) away from the mean was defined as abnormal. For basal GluR level in unstimulated RNAi lines, GluR intensity was compared to control NMJs dissected in parallel, and a p-value < 0.05 compared to control was defined as abnormal. To determine extracellular/secreted lysozyme or neuroglian level, individual synaptic boutons were circled at 0.5 μm beyond HRP staining. Average staining intensity (normalized to bouton area) were always compared to mock controls done within the same experimental set to obtain fold change.

Quantification of GFP-Atg8a, mCherry-Atg8a, and GFP-Lamp1 puncta were performed using Image J. Punctate spots were counted manually and summed from 5 boutons in a continuous Type Ib branch per NMJ, then normalized to the summed area from the same 5 boutons. To ensure that changes in fluorescence signal is not due a difference in protein expression caused by transgene dosage, control NMJs also expressed luciferase in the TRiP RNAi control vector to balance out transgene number. To determine autolysosome number, the number of punctate spots that showed colocalization between mCherry-Atg8a and GFP-Lamp1 were normalized to the total number of autophagic vesicles determined by mCherry-Atg8a signal counted in the same boutons. To determine GFP-Atg8a, mCherry-Atg8a, and GFP-Lamp1 intensities, type Ib boutons were circled based on HRP signal, and the average intensities (normalized to the bouton area) were measured using Image J.

### Kegg pathway, Reactome pathway, and DISEASE analyses

Cytoscape was used to display functional pathway analysis results. The String database plugin was used to retrieve functional enrichment for Gene Ontology terms including cellular component analysis and DISEASE database analysis. Human gene orthologs were used in bioinformatics analysis. KEGG and Reactome analyses were performed using ClueGO plugin in Cytoscape. A minimum of 3 genes in each GO term and a p < 0.05 was used as criteria.

### Statistical analysis

All data are presented as mean ± SEM. Sample numbers are shown in the graphs or figure legends and represent biological replicates. The number of samples used is consistent with established standards in the literature. Samples were randomized during dissection, image collection, and data analyses to minimize bias. To compare unstimulated and stimulated samples of the same genotype, Student’s T-test was used. For multiple samples, One-way ANOVA followed by post hoc analysis with Turkey’s multiple comparison test and was used to determine statistical significance.

## Acknowledgement

We would like to thank all the undergraduate students in the lab over the years for their help with the genetic screen, and members of the lab for their helpful comments and critical reading of the manuscript. K.C. is supported by NIH grants R01NS080946 and R01NS102260.

## Author contributions

Y.C. and Y.G. contributed equally to this work and performed most of the experiments and data analyses. J.Y.L. designed and performed the genetic screen and analyzed data. J.L. performed the genetic screen and contributed to data analysis. K.C. conceived, designed, and supervised all aspects of the project. K.C. wrote the original draft with inputs from Y.C. and Y.G. All authors reviewed and edited the manuscript.

## Competing Interest

The authors declare no competing interests.

## Data Availability

The data that are associated with this manuscript, additional information and requests for reagents used in this study are available upon request.

## List of Supplementary Materials

**Supplementary Figure 1.**
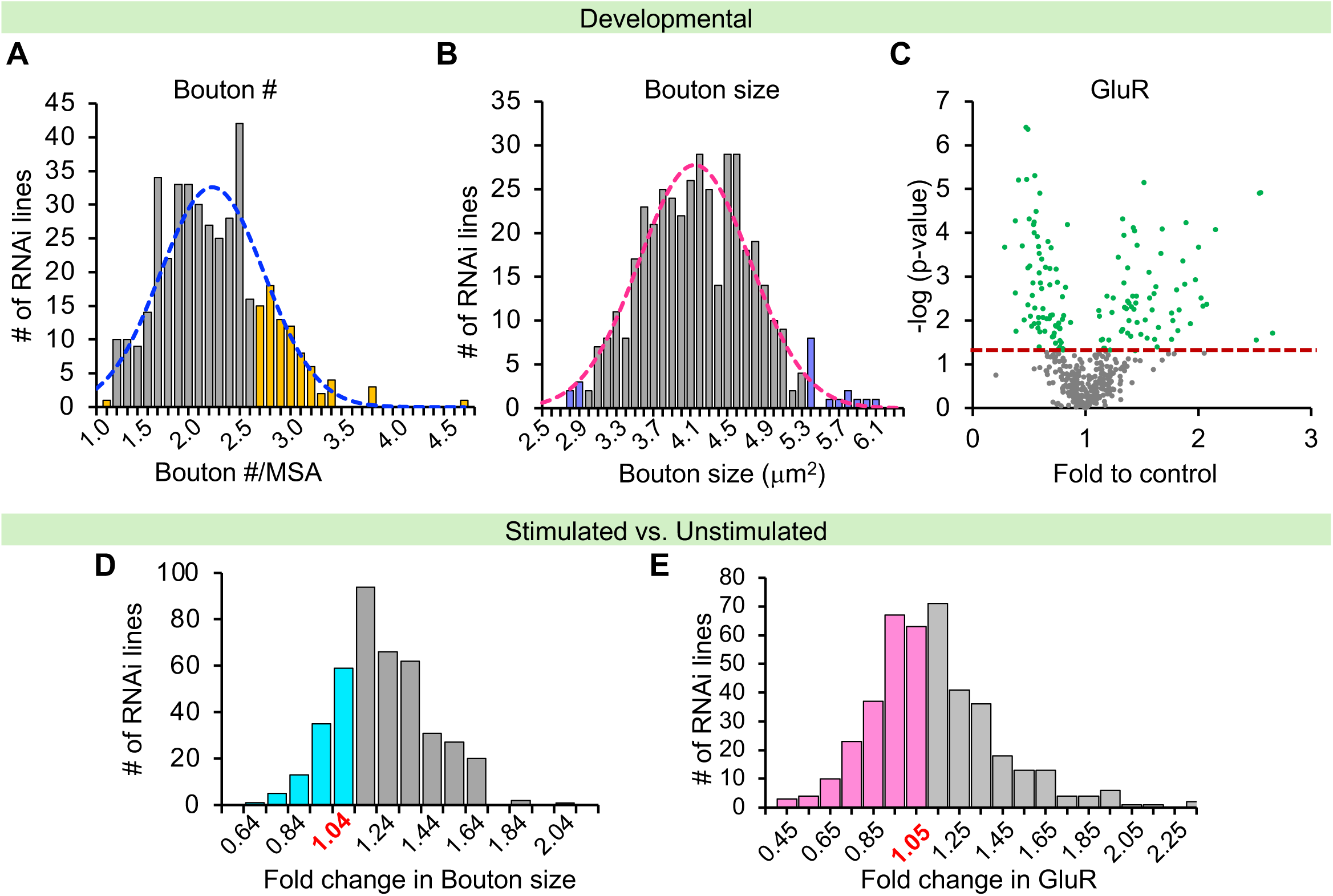
Histogram of the morphological parameters examined in the RNAi-based genetic screen. Histogram for **(A)**, bouton number and **(B)** bouton size. **(C)** Fold change in GluR intensity compared to control and the respective p values obtained for each RNAi line in the screen. Points above the red dashed line represent p-value < 0.05 compared to mock unstimulated control. **(D)** Fold change in activity-induced bouton size. **(E)** GluR intensity. Colored bars highlight the RNAi lines that showed values that are two standard deviations from the average control values.

**Supplementary Figure 2.**
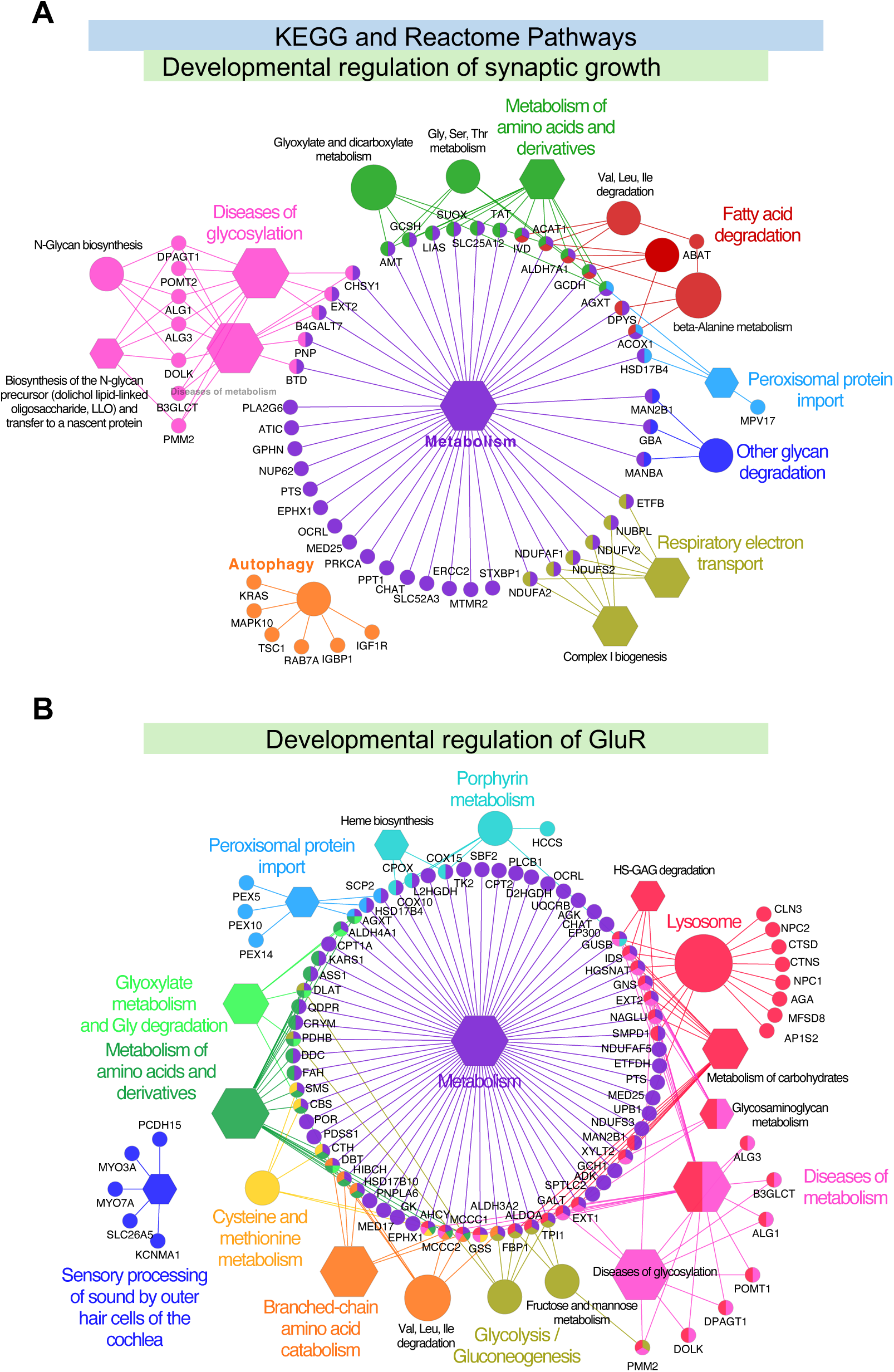
Functional pathway analysis for genes affecting synapse development. KEGG and Reactome pathway analyses for genes affecting **(A)** synaptic growth (bouton number), and **(B)** basal GluR intensity. ClueGo plugin in Cytoscape was used to visualize the networks. KEGG pathway terms are shown in circle and Reactome pathway shown in hexagon. Larger size represents higher significance. Only term description with p-value < 0.05 are shown. Small circles show individual genes (human orthologs) in the functional pathways.

**Supplementary Fig. 3.**
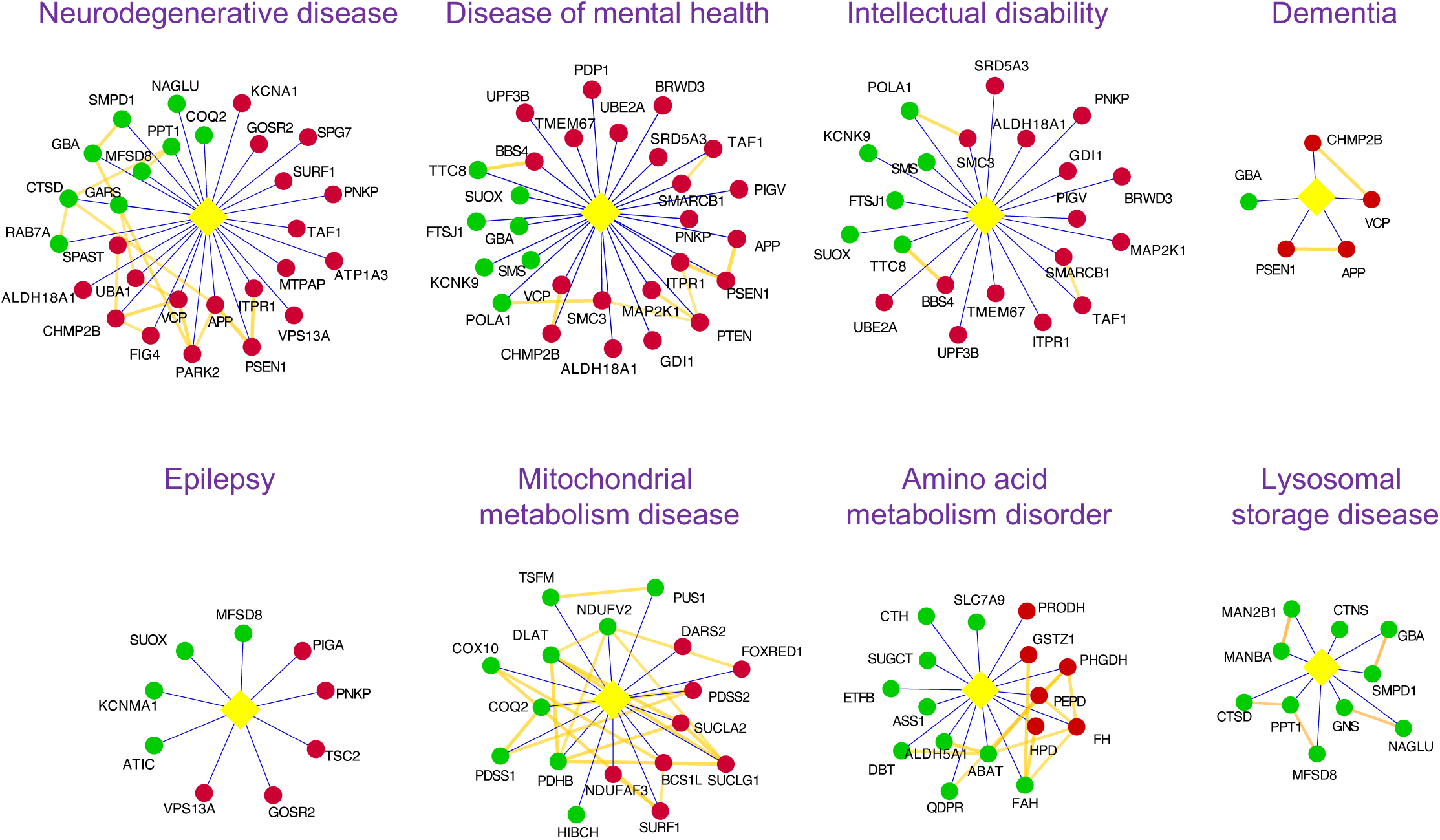
STRING interaction network analysis for genes identified in each disease category using DISEASE database. Red nodes represent Group 1 genes and green nodes represent Group 3 genes. Orange lines indicate high confidence physical interactions identified using STRING.

**Supplementary Figure 4.**
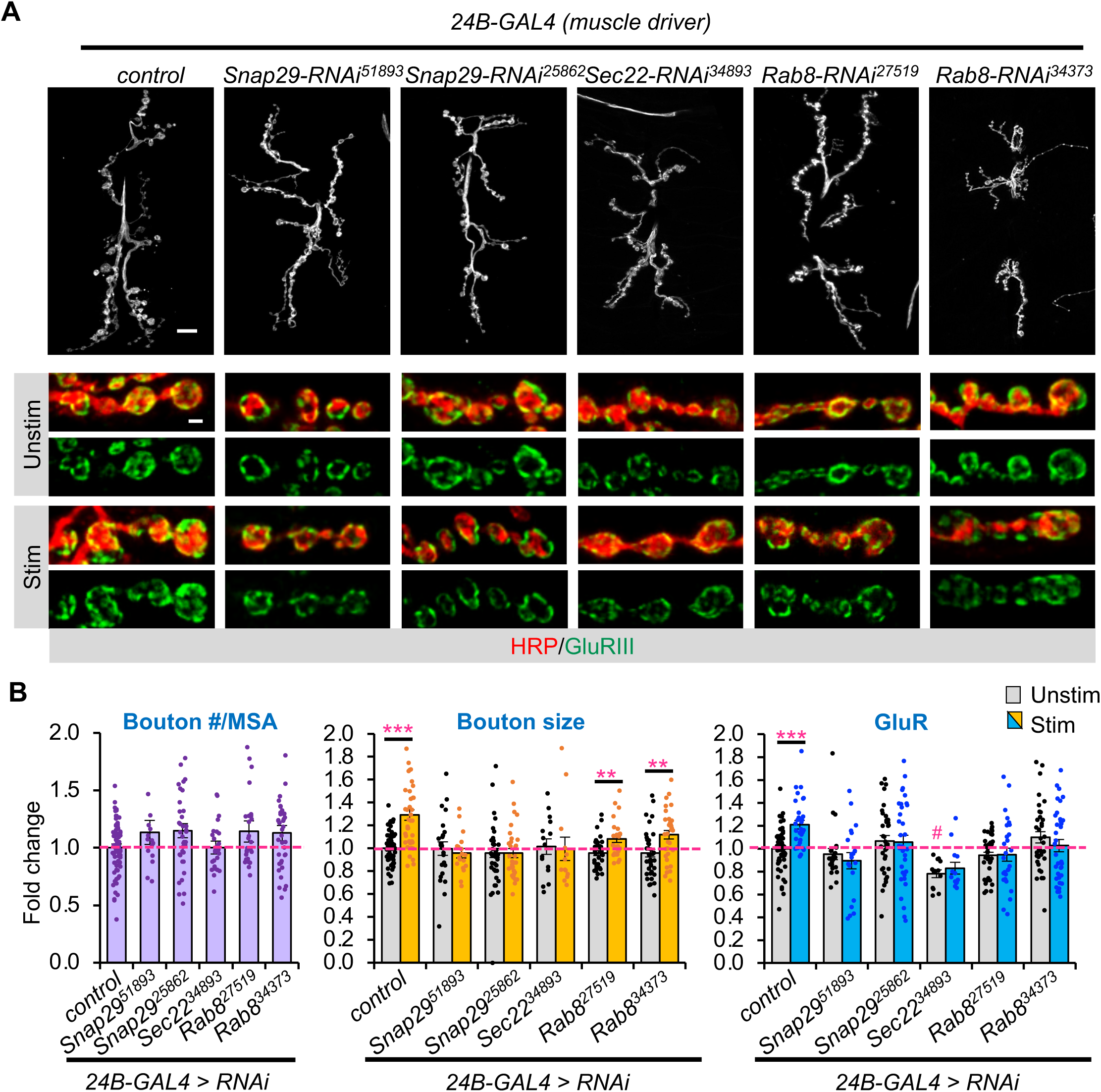
Knockdown of secretory autophagy molecular components in muscles. (A) Gray-scale images of unstimulated NMJs labeled with HRP (top row). Scale bar = 10 μm. Colored images show unstimulated and stimulated synaptic terminals stained with HRP (red) and GluRIII (green). Scale bar = 2 μm. **(B)** Quantification of fold change in bouton number (normalized to muscle surface area), bouton size, and GluR intensity as indicated. Expression in muscles driven by the *24B-GAL4* driver cell-autonomously altered basal GluR level when Sec22 was knocked down. Knockdown of Rab8 in muscles also did not block activity-induced bouton enlargement. All values are mean ± s.e.m. Statistics: One-way Anova followed by Tukey’s multiple comparison test was used to compare between mock control and unstimulated samples across genotypes. Student’s t-test was used to compare between unstimulated and stimulated NMJs of the same genotype. # p ≤ 0.05 when comparing unstimulated samples of different genotypes to unstimulated control. ** p σ; 0.01; *** p σ; 0.001 when comparing stimulated to unstimulated NMJs.

**Supplementary Figure 5.**
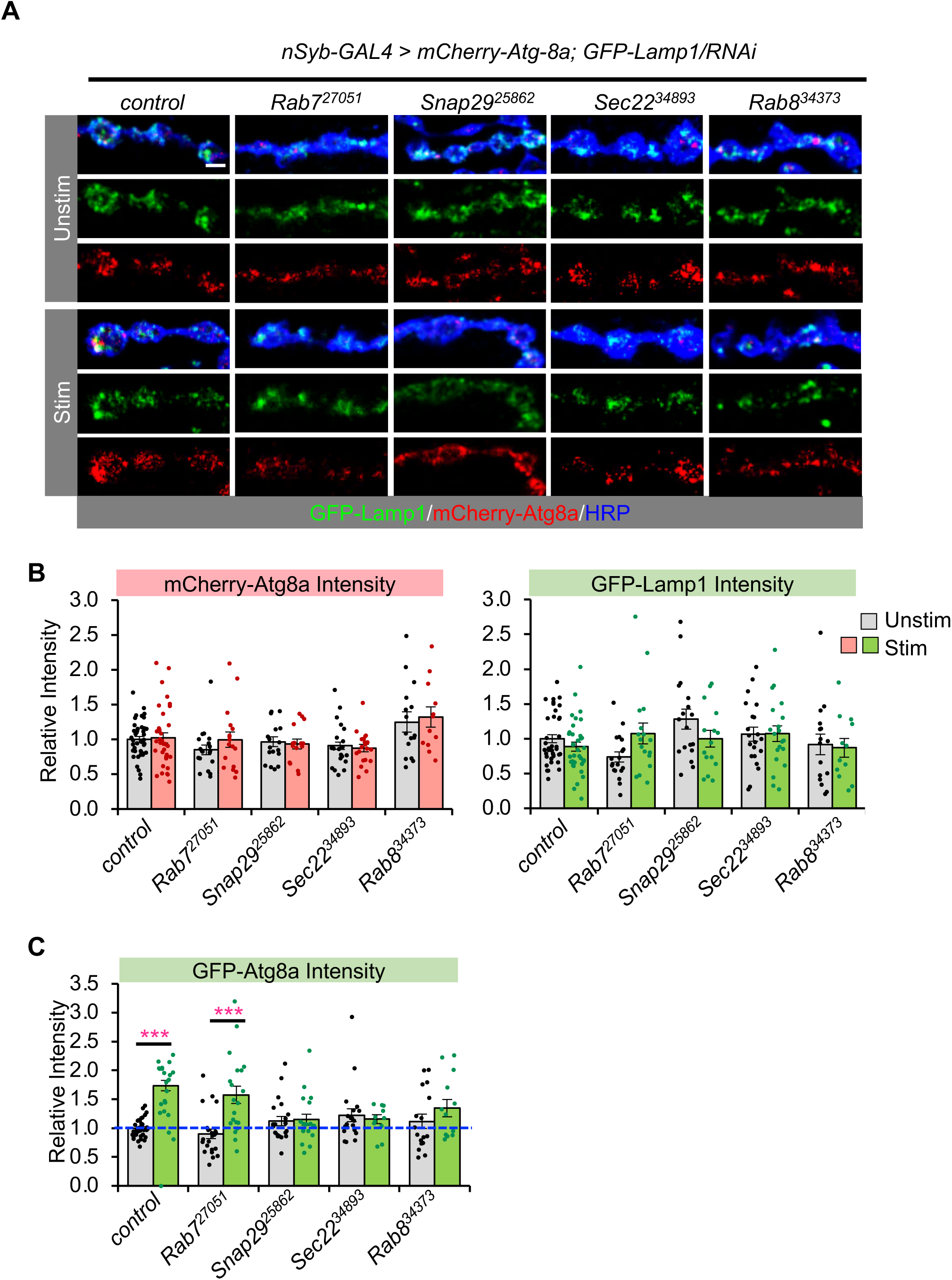
Neuronal activity increases GFP-Atg8a intensity but not mCherry-Atg8a or GFP-Lamp1 intensity. (A) Images of NMJs expressing mCherry-Atg8a, GFP-Lamp1 and the indicated RNAi lines. Scale bar = 2 μm. **(B)** There was no change in the average intensity of mCherry-Atg8a or GFP-Lamp1 at the NMJ. **(C)** Quantification of GFP-Atg8a fluorescence intensity at the NMJ. All values are mean ± s.e.m. Statistics: One-way Anova followed by Tukey’s multiple comparison test was used to compare between mock control and unstimulated samples across genotypes. Student’s t-test was used to compare between unstimulated and stimulated NMJs of the same genotype. *** p σ; 0.001 between stimulated and unstimulated NMJs as indicated.

**Supplementary Figure 6.**
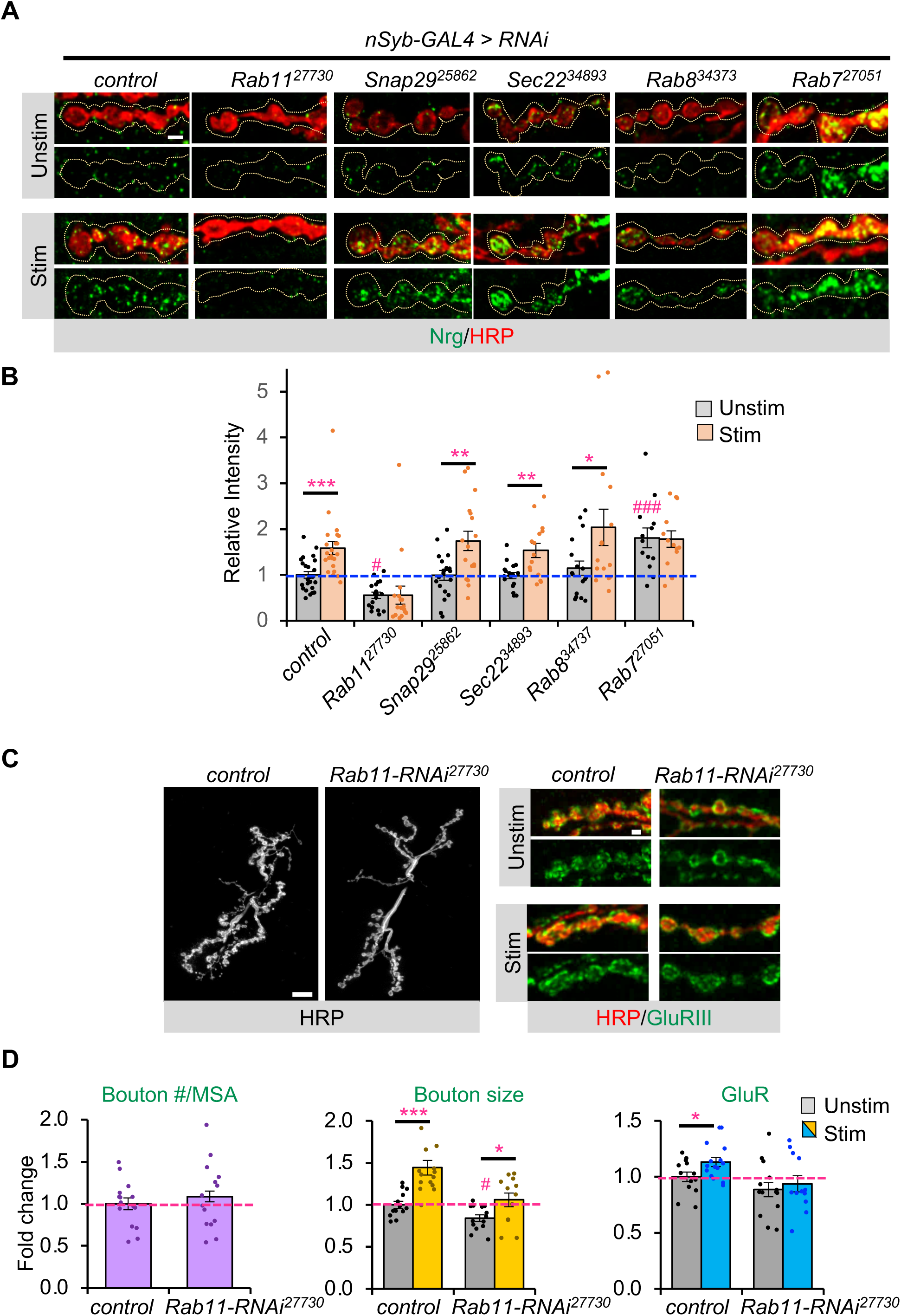
Neuroglian release is through a Rab11-dependent exosomal pathway distinct from secretory autophagy pathway. (A) Images showing extracellular neuroglian (Nrg) staining (green) at the NMJ. The synaptic boutons are outlined by HRP (red). To analyze extracellular Nrg signal, an area that is 0.5 μm beyond the neuronal membrane was circled (dashed line). Scale bar = 2 μm. **(B)** Quantification of extracellular Nrg levels normalized to unstimulated control. Snap29, Sec22, and Rab8, molecular components of secretory autophagy did not block Nrg release. **(C)** Gray-scale images of unstimulated NMJs labeled with HRP. Scale bar = 10 μm. Colored images show unstimulated and stimulated synaptic terminals stained with HRP (red) and GluRIII (green). Scale bar = 2 μm. **(D)** Quantification of fold change in bouton number (normalized to muscle surface area), bouton size, and GluR intensity as indicated. All values are mean ± s.e.m. One-way Anova followed by Tukey’s multiple comparison test was used to compare between mock control and unstimulated samples across genotypes. Student’s t-test was used to compare between unstimulated and stimulated NMJs of the same genotype. # p ≤ 0.05; ### p ≤ 0.001 when compared to unstimulated control. * p σ; 0.05; ** σ; 0.01; p *** p σ; 0.001 between stimulated and unstimulated NMJs of the same genotype.

**Data S1.** List of RNAi lines used in the genetic screen.

**Data S2.** Group classification for each gene in the genetic screen.

## Notes

### Competing Interest Statement

The authors have declared no competing interest.

